# Unbiased and Epicardial-Specific Lineage Tracing Reveal Epicardial Contribution to Vascular Endothelial Cells in Heart Development

**DOI:** 10.64898/2026.08.23.742225

**Authors:** Payal Ghosh, Zibei Gao, Haoting He, Juan Xu, Guang Li

## Abstract

The lineage potential of cardiac cells, particularly epicardial cells, during heart development remains unclear, largely due to the non-specific expression of epicardial marker genes and the resulting non-specific labeling in Cre-loxP mouse models. Using DARLIN mice, a CRISPR/Cas9-based lineage-tracing system independent of the Cre-loxP system, we analyzed the lineage development of embryonic cardiac cells in an unbiased manner and identified lineages shared among different cell types, such as epicardial cells and vascular endothelial cells (Vas_ECs). To further confirm the lineage potential of epicardial cells, we identified an epicardial cell-specific marker gene, Lrrn4, through analysis of a multi-staged single-cell mRNA-sequencing (scRNA-seq) dataset, and generated a corresponding Lrrn4-CreER mouse line. We then bred this line with a reporter mouse to confirm its specificity for labeling epicardial cells, and subsequently performed prolonged lineage tracing, which revealed differentiation of the labeled epicardial cells into Vas_ECs and fibroblasts, but not cardiomyocytes, indicating that Lrrn4 labels a population of Epi with the potential to differentiate into Vas_ECs. Finally, Using this mouse line, we investigated epicardial cell function by selectively ablating these cells and by expressing TGFβ in epicardial cells to convert their lineage from Vas_ECs to fibroblasts. Both approaches resulted in significant developmental defects in embryonic hearts. Together, these results indicate that epicardial cells can give rise to Vas_ECs, and that the Lrrn4-CreER mouse model is a valuable tool for elucidating the role of the epicardium in heart development.

## Introduction

Epicardial cells form a single cellular layer covering the outer surface of the heart. They function as progenitors capable of differentiating into multiple cardiac cell types, in addition to serving as a signaling hub that regulates the development of other cardiac cell lineages^1–5^. Epicardial cells are derived from juxta-cardiac field (JCF) progenitors at the cardiac crescent stage, which have been reported to give rise to both epicardial cells and cardiomyocytes^6–8^. Afterward, the progenitors were reported to derive from a transient tissue structure known as the proepicardial organ (PEO)^9^, which is known to be heterogeneous and consists of pro-epicardial (PE) cells, vascular smooth muscle cell progenitors, and sinoatrial progenitors^10^. The contribution of PE cells to epicardial cells was largely completed at early embryonic stage E10.5^11^.

Epicardial cells have been reported to highly express multiple genes, such as Wt1, Tbx18, Aldh1a2, and Krt19^12,13^. These genes have been used to generate multiple transgenic mouse lines, and lineage-tracing studies using these lines have revealed distinct lineage descendants ^15–18^, suggesting that epicardial cells may represent a heterogeneous population. However, these genes have also been reported to be expressed in other cardiac cell types besides the epicardium. For instance, Wt1 is also expressed in endothelial cells, fibroblasts, and cardiomyocytes, while Tbx18 is expressed in fibroblasts and smooth muscle cells^14^. This expression in non-epicardial cell types complicates the interpretation of lineage-tracing results based on these markers, raising questions about the true lineage potential of the epicardial cells they label.

DARLIN is a CRISPR/Cas9-based lineage-tracing mouse strain that reconstructs lineage relationships based on the history of accumulated genetic edits. It consists of three Cas9-targeting arrays, named CA, TA, and RA, with the theoretical capacity to generate up to 10^18^ distinct barcodes^19,20^. In addition, Cas9 expression is controlled by a Tet-On system, allowing genetic editing and lineage tracing to be initiated at a defined time point through the conditional addition of doxycycline (Fig. S1). This system has been successfully applied to studying the development of hematopoietic lineages in both embryonic and adult mice^19,20^. Because this approach does not rely on the traditional Cre-loxP system to trace lineages, it avoids the non-specific expression and labeling associated with conventional lineage-tracing mouse lines.

Furthermore, through analysis of scRNA-seq data spanning embryonic and neonatal stages^14^, we identified leucine-rich repeat neuronal protein 4 (Lrrn4) as being specifically expressed in epicardial cells across these stages. Lrrn4 is a membrane-associated protein containing leucine-rich repeat and fibronectin type III domains^21,22^. It has been implicated in long-term memory and spatial learning, while Lrrn4 mutants otherwise undergo normal embryonic development^22^. Given its epicardial cell-specific expression pattern, the Lrrn4 locus represents an ideal site for specifically labeling this cell type. To this end, we inserted a CreER sequence immediately downstream of the Lrrn4 coding sequence to generate a mouse line named “Lrrn4-CreER.”

In this study, by combining unbiased lineage tracing using DARLIN mice with epicardial-specific lineage tracing using the newly generated Lrrn4-CreER mouse line, we found that the epicardial cell lineage contributes to multiple cell lineages, including Vas_EC. We further overexpressed TGFβ in epicardial cells to redirect their lineages, and separately bred the Lrrn4-CreER line with a DTA mouse to selectively ablate epicardial cells, revealing an important role for epicardial cells in heart development.

## Results

### DARLIN lineage tracing

Following four-chamber formation at E10.5, mouse embryonic hearts begin diversifying into distinct lineages, such as fibroblasts and vascular endothelial cells (Vas_ECs), starting around E12.5. To understand lineage specification during this time window, we treated pregnant DARLIN mice with doxycycline at E10.5 to induce Cas9 expression and harvested the treated embryonic hearts at E14.5 (Fig. 1A, S1A). We then sequenced the CRISPR-generated edits and the transcriptional profile of single cells within the hearts. Based on the expression of lineage-specific genes in the scRNA-seq dataset, we identified all major cardiac cell types, including atrial cardiomyocytes (Atrial_CM), ventricular cardiomyocytes (Ven_CM), endocardial endothelial cells (Endo_EC), vascular endothelial cells (Vas_EC), epicardial cells (Epi), smooth muscle cells (SMC), pericytes, fibroblasts (Fb), and immune cells (Fig. 1B, C). Among these cell types, Ven_CM, Fb, and Endo_EC were the most abundant (Fig. 1D).

**Fig. 1.**
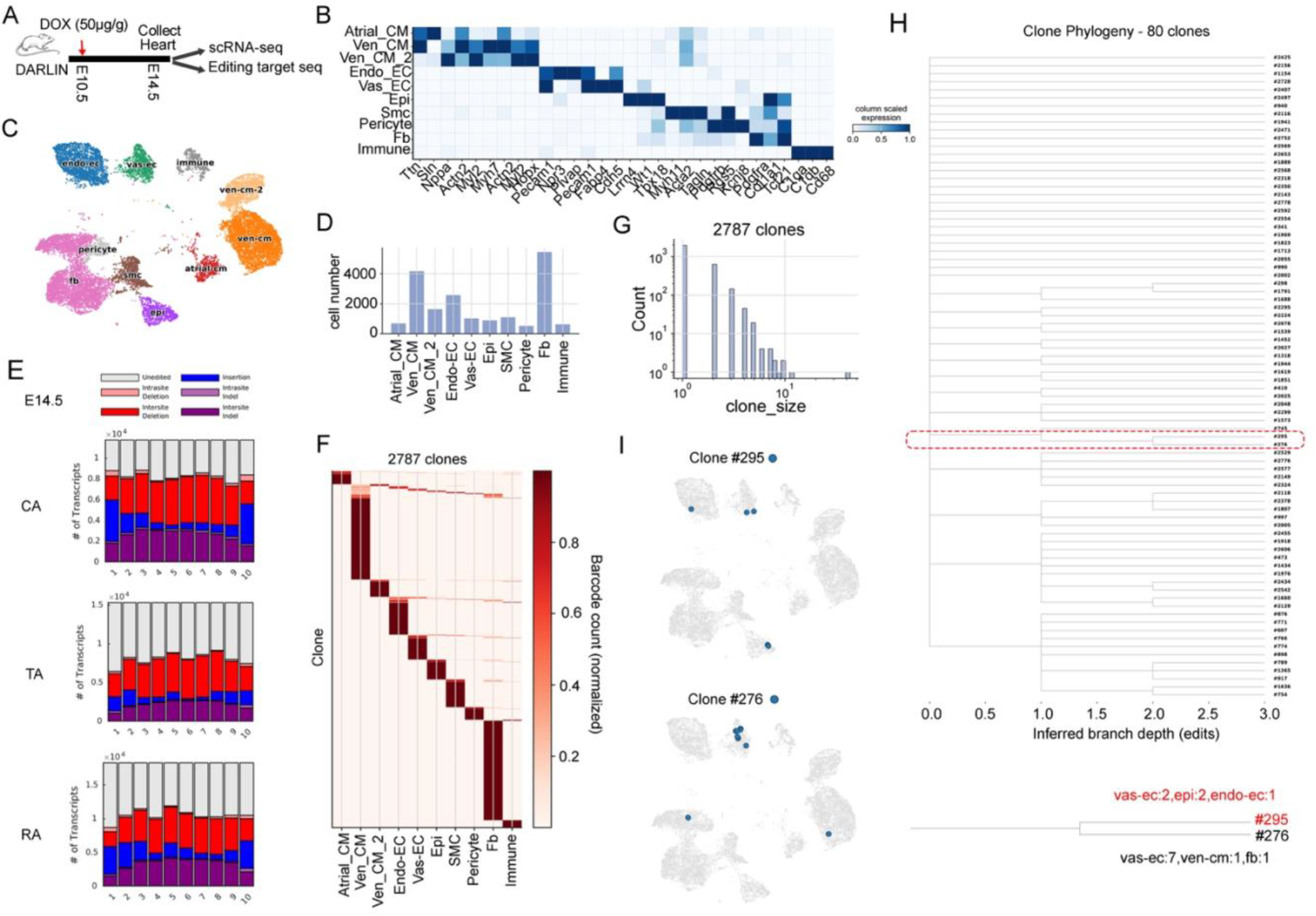
Lineage analysis of mouse embryonic heart development at E14.5 using DARLIN. (A) Diagram of the lineage tracing experiment with Dox administration at E10.5 and heart collection at E14.5. (B) Heatmap of the lineage genes expression in each cell type. (C) UMAP plot of the scRNA-seq data labeled with cell type information. (D) The number of cells recovered at E14.5 scRNA-seq dataset. (E) UMI factions on each targeting sites. (F) Heatmap showing the cellular compositions within each clone in the E14.5 sample. (G) Histogram showing the number of clones at each clone size. (H) Phylogeny of the clones at E14.5. (I) UMAP plots of two representative clones, with cell types in each clone labeled.

Next, we analyzed the CRISPR editing sequences and recovered various types of edits across the editing sites distributed across the three targeting arrays, CA, TA, and RA (Fig. 1E, S1B). By integrating the CRISPR editing information with the scRNA-seq profiles, we successfully identified 2,787 cell clones, most of which consisted of a single lineage (Fig. 1F). The number of clones decreased with increasing clone size overall, with most recovered clones containing fewer than 10 cells (Fig. 1G). We additionally identified 80 clones containing multiple cell lineages. Among these, 13 clones contained Epi together with another lineage: 6 clones consisted of Epi and Fb, 3 clones consisted of Epi and Ven_CM, 3 clones consisted of Epi and Vas_EC, and 1 clone consisted of Epi and Endo_EC. The remaining 67 clones represented other multi-lineage combinations not involving Epi (Table S1). The presence of multiple cell lineages within a single clone suggests that these clones arose from a common progenitor capable of giving rise to several cell types. Because lineage tracing was initiated at E10.5, by which point the epicardial layer has largely formed, these progenitors were most likely epicardial cells themselves, indicating that the epicardium is a multipotent and heterogeneous cell population. We further constructed a phylogenetic tree for the 80 clones and identified branches containing multiple related clones. For example, clones 295 and 276 shared a common branch: clone 295 consisted of 2 Epi, 2 Vas_EC, and 1 Endo_EC, while clone 276 consisted of 7 Vas_EC, 1 Ven_CM, and 1 Fb. These results suggest that the progenitors of these two clones had the potential to differentiate into multiple cell types, including Epi and Vas_ECs (Fig. 1H, I).

We next used the same system to analyze lineage development over a longer developmental window. Doxycycline was again administered at E10.5, and embryonic hearts were collected at E18.5 to analyze the transcriptional profile and lineage-associated edits of each single cell (Fig. S2A). Based on lineage gene expression, we identified the cell types present in the scRNA-seq data and found that Fb and Vas_EC were the two most abundant cell types (Fig. S2B–D). We then integrated the scRNA-seq data with the lineage-tracing edits and successfully recovered 4,187 clones (Fig. S2E, F). Overall clone sizes were slightly smaller than those observed at E14.5 (Fig. S2G), likely reflecting the greater number of cells and clones generated by E18.5 relative to E14.5, given that a similar number of cells were targeted at the two stages. Among these clones, 71 consisted of multiple cell lineages, while the remainder contained only a single lineage. Of these, 22 clones contained Epi together with another lineage: 11 clones consisted of Epi and Fb, 4 clones consisted of Epi and Vas_EC, 2 clones consisted of Epi and atrial CM, 2 clones consisted of Epi, Fb, and immune cells, 1 clone consisted of Epi and SMC, 1 clone consisted of Epi and pericyte, and 1 clone consisted of Epi and Ven_CM. The remaining 49 clones represented other multi-lineage combinations not involving Epi (Table S2). We further reconstructed the phylogenetic tree for these clones and identified branches containing multiple related clones (Fig. S2H, I). For example, clone 3132, which consisted of 1 Epi, 1 Ven_CM, and 1 Vas_EC, shared a branch with clones 3145, 910, and 1934. Together, these analyses suggest that Epi may share a common developmental path with multiple distinct cell types, including Vas_EC.

### Lrrn4 is specifically expressed in epicardial cells

Through analysis of scRNA-seq data from 18 developmental stages spanning embryonic and neonatal hearts, we found that Lrrn4 is specifically expressed in epicardial cells across all stages examined. Although Lrrn4 was expressed in most epicardial cells, it did not cover every single cell (Fig. 2A, B). We further analyzed single-cell multiome sequencing data from E14.5^23^ embryonic hearts and observed high Lrrn4 RNA expression in epicardial cells. We also found that ATAC-seq peaks surrounding the Lrrn4 genomic locus were specifically enriched in epicardial cells, with regulatory interactions being detected (Fig. 2C). Analysis of scATAC-seq data from P4 cardiac cells similarly showed that ATAC-seq peaks at the Lrrn4 locus were specifically enriched in epicardial cells (Fig. S3A).

**Fig. 2.**
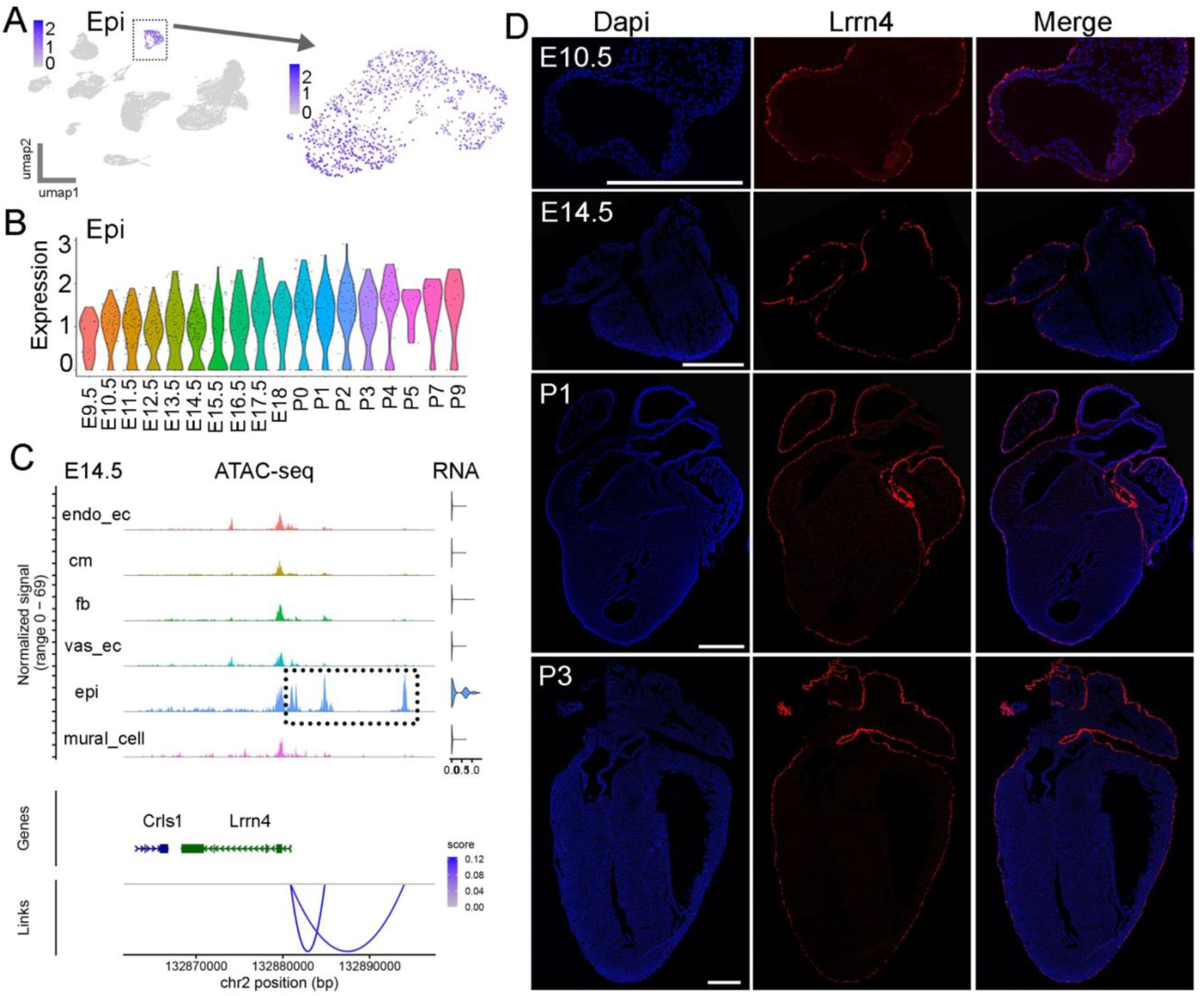
Lrrn4 is specifically expressed in epicardial cells. (A) UMAP plot of scRNA-seq data showing epicardial-specific Lrrn4 expression. (B) Violin plot of Lrrn4 expression in epicardial cells across 18 developmental stages. (C) Single-cell multiome-seq data at E14.5 showing epicardial-specific Lrrn4 expression and enriched ATAC-seq peaks at its genomic locus. (D) RNA staining showing epicardial-specific Lrrn4 expression at multiple stages. Scale bar = 500 μm.

Next, we examined Lrrn4 expression using RNA staining and observed epicardial cell-specific expression across multiple stages, including E10.5, E14.5, P1, and P3 (Fig. 2D). Interestingly, Lrrn4 was not uniformly expressed in all cells of the epicardial layer. Instead, clear Lrrn4-negative cells were interspersed among Lrrn4-positive cells at each stage, and this pattern was largely consistent across all four chambers (Fig. 2D). To further characterize this heterogeneity, we compared the single-cell transcriptional expression of Lrrn4, Wt1, Tbx18, and Krt19, revealing both shared and distinct expression patterns among epicardial cells at different stages (Fig. S4). We next performed co-staining of Lrrn4 with Tbx18 and Wt1 at E14.5 and P3. Lrrn4 expression remained restricted to epicardial cells at both stages, whereas Wt1 and Tbx18 were also detected within the heart chambers, beyond the epicardium (Fig. S5A–D). Within the epicardial cell population itself, Lrrn4 again showed both shared and distinct expression relative to the other two genes (Fig. S5A–D). Together, these results indicate that Lrrn4 is specifically expressed in epicardial cells, comprising a mixture of cells co-labeled by other epicardial marker genes and cells uniquely labeled by Lrrn4 alone.

### Lrrn4-CreER mice label epicardial cells

To label Lrrn4-positive cells, we generated a transgenic mouse line by inserting a CreER coding sequence at the 3′ end of the Lrrn4 genomic locus using CRISPR/Cas9. The CreER coding sequence was linked to the final exon of Lrrn4 via a P2A sequence (Fig. 3A). We then crossed Lrrn4-CreER mice with the reporter line Rosa26-mTmG. Pregnant mice were treated with tamoxifen from E9.5 to E11.5, and embryos were collected at E12.5 (Fig. 3B). Using eGFP imaging and ESR staining of heart sections, we found that both signals were specifically localized to epicardial cells (Fig. 3C). Further staining for Aldh1a2 revealed substantial overlap between eGFP and Aldh1a2 signals on the heart surface (Fig. 3D), confirming that the labeled cells were epicardial cells.

**Fig. 3.**
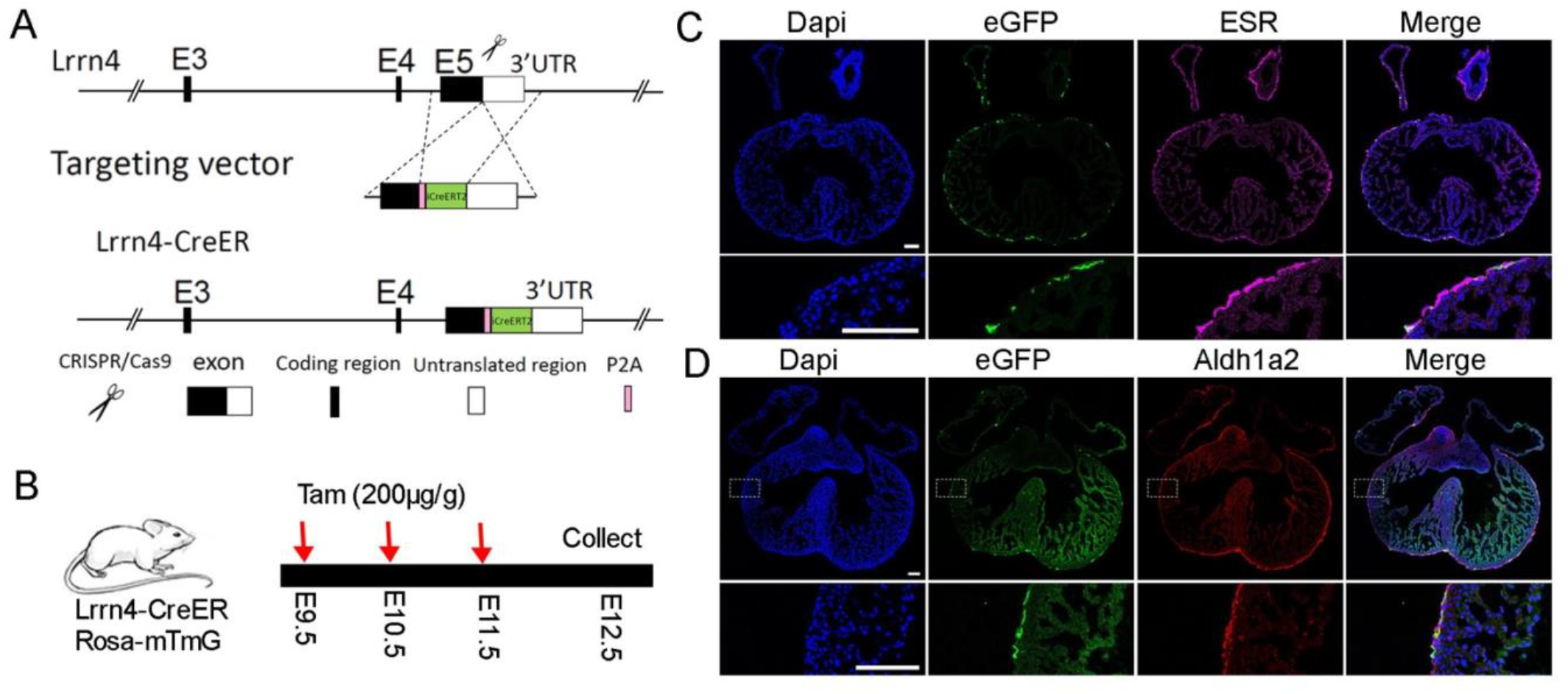
Lrrn4-CreER; Rosa26-mTmG specifically labels epicardial cells. (A) Schematic of the strategy used to generate Lrrn4-CreER transgenic mice. (B) Lineage labeling strategy by crossing Lrrn4-CreER mice with Rosa26-mTmG mice. (C) eGFP and ESR signals are specifically enriched in epicardial cells. (D) eGFP signal overlaps extensively with Aldh1a2 expression. Scale bar = 100 μm.

### Lineage tracing analysis of Lrrn4-CreER–labeled cells

We next examined the lineage contribution of Lrrn4-CreER–labeled cells by analyzing hearts several days after tamoxifen treatment. Pregnant mice were treated with tamoxifen from E9.5 to E11.5, and hearts were collected at E17.5 (Fig. 4A). We first confirmed that eGFP-positive cells on the heart surface were Wt1-positive (Fig. 4B). Staining for Pdgfra revealed that a substantial proportion of eGFP-positive cells were also Pdgfra-positive (Fig. 4C), indicating differentiation into fibroblasts. In addition, staining for Cd31 identified many eGFP/Cd31 double-positive cells (Fig. 4D), suggesting that Lrrn4-labeled epicardial cells also contributed to endothelial cells.

**Fig. 4.**
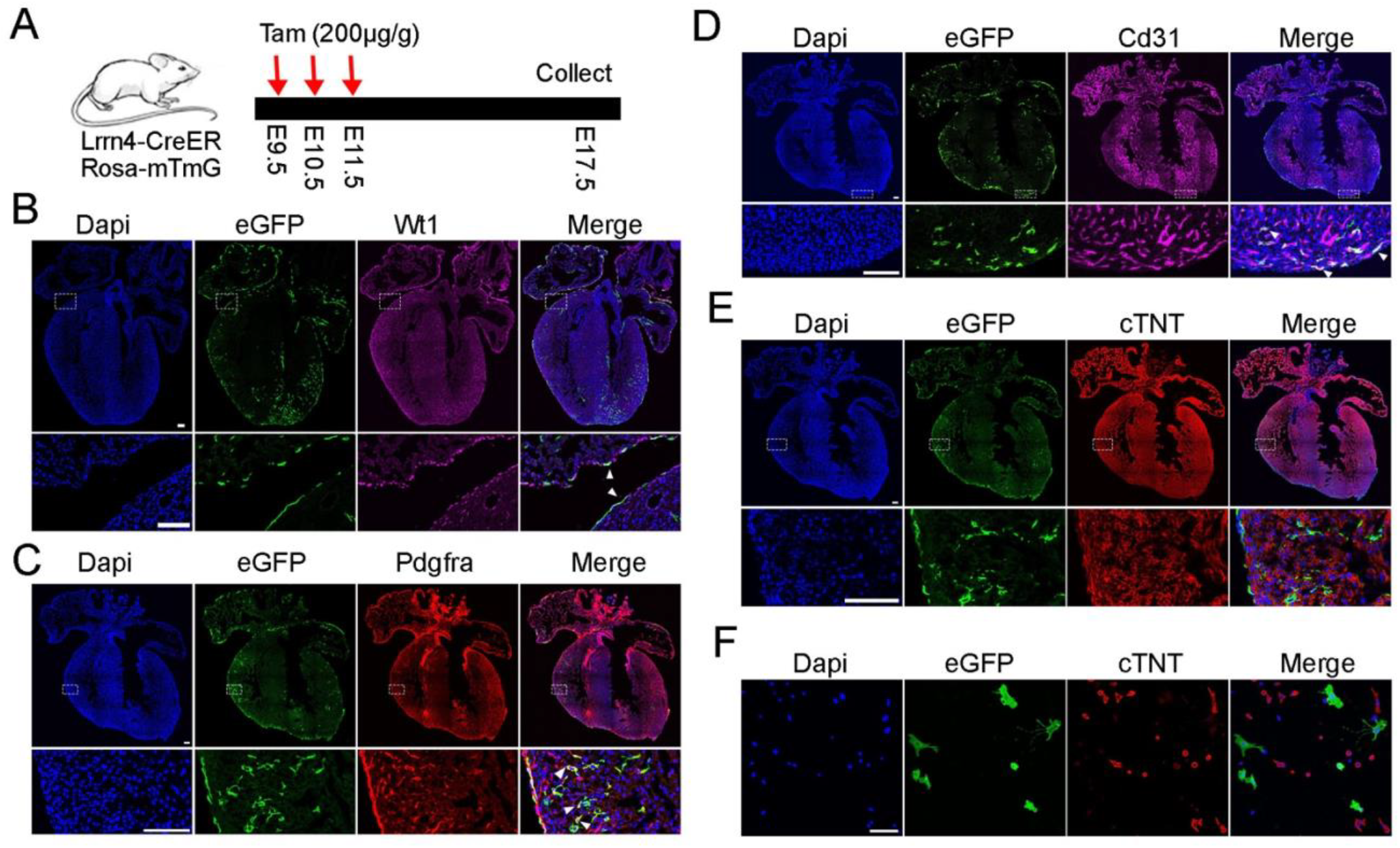
Lineage tracing analysis of Lrrn4-CreER–labeled cells. (A) Experimental design for lineage tracing. (B) Wt1 staining confirms that eGFP-positive cells on the heart surface are epicardial. (C) Pdgfra staining identifies fibroblast differentiation of eGFP-positive cells. (D) Cd31 staining identifies endothelial differentiation of eGFP-positive cells. (E, F) No cTNT-positive cardiomyocytes overlap with eGFP signals in tissue sections or cultured cells. Scale bar = 100 μm.

In contrast, staining for cTNT showed no overlap with eGFP in heart sections (Fig. 4E). We further dissociated cardiac cells and cultured them overnight, with or without FACS enrichment for eGFP-positive cells. We did not observe clear eGFP/cTNT double-positive cells, aside from one potential candidate cell (Fig. 4F, S6A). Similar experiments were performed by administering tamoxifen at E10.5 and E11.5, followed by staining for Aldh1a2, Pdgfra, Cd31, and cTNT, and yielded consistent results (Fig. S7A, B). Collectively, these findings indicate that Lrrn4-CreER–labeled epicardial cells contribute to fibroblast and endothelial cell lineages, but likely not to cardiomyocytes.

### Functional analysis of epicardial cells

Subsequently, we analyzed the function of the epicardium by breeding Lrrn4-CreER mice with Rosa26-DTA mice to specifically ablate epicardial cells. Pregnant mice were treated with tamoxifen at E10.5 and E11.5, and embryos were collected at E14.5 (Fig. 5A). Control embryos and embryonic hearts appeared normal, whereas the ablated embryos appeared to be dead, and the corresponding embryonic hearts also appeared to have ceased further development (Fig. 5B), indicating the importance of epicardial cells in embryonic heart and overall embryo development.

**Fig. 5.**
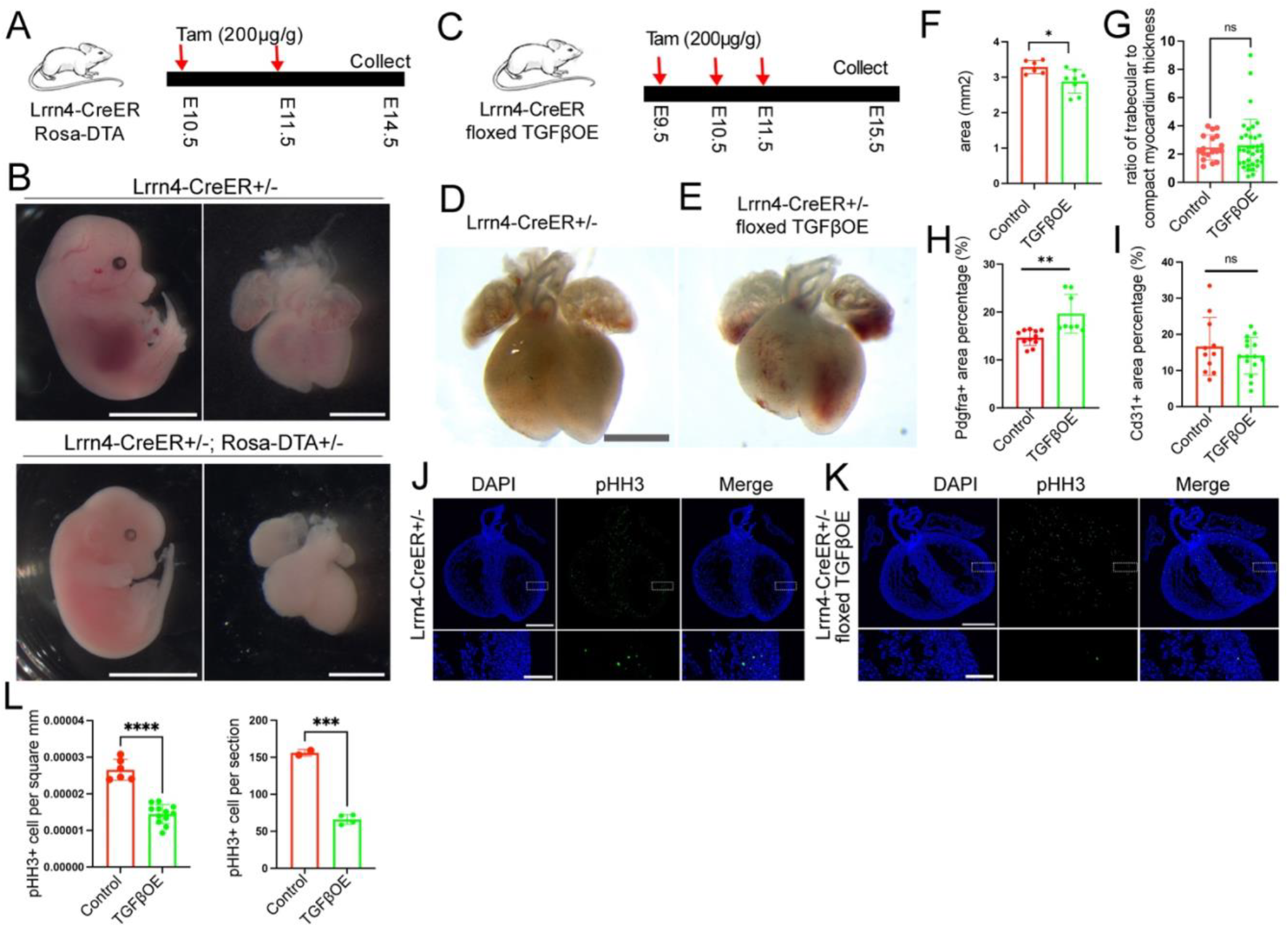
Functional analysis of epicardium through ablation and TGFβ1 ectopic expression. (A) Diagram of the experiment to ablate epicardial cells. (B) Ablation of Lrrn4+ cells led to embryonic lethality. (C) Diagram of the strategy to overexpress TGFβ1 in epicardial cells. (D-F) The epicardial cell-specific TGFβ1 overexpression led to smaller hearts. (G) Quantification of the ratio of trabecular to compact myocardium thickness. (H, I) Quantification of fibroblast and endothelial cell areas in the left ventricle of control and TGFβ1-overexpression mice. (J, K) Representative images of control and TGFβ1 overexpression embryonic hearts stained with antibody for pHH3. (L) Quantification of pHH3 positive cells per section and area. Data are presented as mean ± SD. *p < 0.05, **p < 0.01, ***p < 0.001, ****p < 0.0001. ns, not significant.

Next, we modulated the lineage choices of Epi cells by specifically expressing TGFβ1 in Lrrn4 positive cells, since TGFβ is known to promote epithelial-to-mesenchymal transition (EMT). By breeding Lrrn4-CreER mice with a floxed TGFβ1-overexpression (βglo) line^24^, we aimed to overexpress TGFβ1 in the epicardium and thereby convert the lineage fate of these cells from Vas_ECs to fibroblasts. Pregnant mice were treated with tamoxifen from E9.5 to E11.5, and embryonic hearts were harvested at E15.5 (Fig. 5C). TGFβ1-overexpressing hearts were found to have reduced size (Fig. 5D–F), and further IF staining revealed an increase in Pdgfra positive fibroblast area relative to controls, along with a slight decrease in Cd31 positive endothelial cell area (Fig. 5H, I).

Given that TGFβ is also known to regulate cell proliferation, we next examined its effect on myocardial growth. During normal development, TGFβ1 is mainly expressed in endocardial ECs (Endo_EC) and restricts CM proliferation in the adjacent trabecular myocardium^25^, whereas the compact myocardium, which lies farther from Endo_EC, exhibits higher CM proliferation, hypothesized to be driven primarily by epicardium-derived signals. To test this model, we performed IF staining for pHH3 and found reduced proliferation in overexpression hearts relative to controls (Fig. 5J–L). Although quantification of the trabecular-to-compact myocardium thickness ratio revealed no difference between control and overexpression hearts (Fig. 5G), the reduced heart size and pHH3 signal together suggest that the epicardium plays an important role in embryonic heart development by promoting compact myocardium growth.

## Discussion

In this study, we used DARLIN mice to unbiasedly analyze cardiac lineage development and identified clones consisting of both epicardial cells and Vas_ECs. We further demonstrated that the Lrrn4-CreER mouse line specifically labels epicardial cells, and that epicardial cells can contribute to multiple cell lineages, including Vas_ECs. Finally, we used this mouse line to investigate the role of epicardial cells in heart development.

CRISPR editing in the DARLIN system is temporally controlled by a Tet-On system, which is activated upon administration of doxycycline^19^. Additionally, because Cas9 and sgRNA expression are driven by ubiquitous promoters, the lineage-tracing results generated by this system are largely free of the bias introduced by leaky expression from cell type-specific promoters in conventional approaches. In this study, we administered doxycycline at E10.5, by which point epicardial cells have emerged from the proepicardial organ (PEO) and covered the heart surface^11^, making this an ideal stage for studying epicardial cell lineage specification. However, studying other aspects of lineage diversification, such as the differentiation of cardiac progenitor cells, would require inducing the system at an earlier or later stage.

Although we did not observe differentiation of Lrrn4-CreER-labeled epicardial cells into cardiomyocytes, this does not conclusively rule out the potential of epicardial cells to give rise to cardiomyocytes. Because Lrrn4-CreER does not label the entire population of epicardial cells, the remaining, unlabeled epicardial cells may still retain this developmental potential. Future studies comparing the transcriptional profiles and lineage contributions of Lrrn4-positive versus Lrrn4-negative epicardial cells would be informative in this regard. Additionally, we observed that some clones identified through DARLIN lineage tracing contained both epicardial cells and ventricular cardiomyocytes (Ven_CMs), raising the possibility that these two lineages share a common progenitor.

In contrast to existing epicardial lineage-tracing lines, which typically begin labeling epicardium-derived cells (EPDCs) only at late embryonic stages^26^, Lrrn4 exhibits epicardial cell-specific expression throughout both embryonic and neonatal stages. This property makes the Lrrn4-CreER mouse line a valuable tool for studying epicardial lineage contributions, epicardial cell function, and gene regulation across all developmental stages, including the late embryonic and neonatal periods.

The epicardium is anatomically adjacent to the compact myocardium, whereas the endocardium lies in close proximity to the trabecular myocardium. Both tissues are known to serve as hubs for growth factor signaling that instructs heart morphogenesis^27,28^. Beyond differences in cardiomyocyte proliferation, the compact and trabecular myocardium also differ in their maturation and function^29^. It would therefore be of interest to examine the roles of other growth factors that are differentially expressed between the epicardium and endocardium. In this context, the Lrrn4-CreER mouse line offers a valuable tool for investigating the function of endocardium-derived factors by enabling their enforced expression in the epicardium.

## Materials and methods

### Mouse lines

The animal experiments were approved by the University of Pittsburgh Institutional Animal Care and Use Committee (IACUC). The mouse lines CA/TA/RA (Strain #038750)^20^, Cas9-TdT-gRNAs-M2 (Strain #038749)^20^, mTmG (Strain #007676)^30^, ROSA26-DTA (Strain #032087)^31^, and β1glo (Strain #018393)^24^ were ordered from the Jackson Laboratory.

### Mouse breeding

The DARLIN mouse model was generated by breeding CA/TA/RA mice with Cas9-TdT-gRNAs-M2 mice. Pregnancies of the female mice were timed. 50 μg of Doxycycline per gram of mouse body weight was administered at E10.5, and embryos were harvested at E14.5 or E18.5. Lrrn4-CreER mice were bred with Rosa26-mTmG mice, and 200 μg of tamoxifen per gram of mouse body weight was administered at specific pregnancy stages ranging from E9.5 to E11.5. Embryonic hearts were collected at E12.5 or E17.5 to analyze the eGFP pattern and for IF staining of ESR or lineage gene expression. Lrrn4-CreER mice were bred with Rosa26-DTA mice, and 200 μg/g of tamoxifen was administered at E10.5 and E11.5, and embryos were collected at E14.5. Lrrn4-CreER mice were also bred with β1glo mice, and tamoxifen was administered from E9.5 to E11.5, and embryonic hearts were collected at E15.5.

### DARLIN mice: scRNA-seq and lineage barcode libraries preparation

Mouse embryonic hearts at E14.5 or E18.5 were dissected and dissociated into single cells through digestion with 0.25% trypsin and a collagenase A/B mixture. The cell concentration was then adjusted to target roughly 25,000 cells per 10x well. This adjusted suspension was loaded onto the Chromium chip for GEM generation and reverse transcription into cDNA. Following cDNA amplification and cleanup, the endogenous transcript library was constructed per the 10x Genomics Chromium Single Cell 3’ Reagent Kit user guide (CG000315 Rev E). Ten microliters of purified cDNA was used for Fragmentation, End Repair & A-tailing. This was followed by a double-sided SPRIselect size selection, Adaptor Ligation, a post-ligation SPRIselect cleanup, and Sample Index PCR using the Dual Index Plate TT Set A to add sample-specific indices. The libraries were then QC’d and sequenced.

The DARLIN lineage-barcode library was built from the same single-cell cDNA generated during the 10x Chromium 3’ GEX workflow^19^. Five microliters of the cDNA were used in three separate first-round PCR reactions (CA, TA, and RA, corresponding to the three DARLIN array types), each containing Kapa HiFi HotStart ReadyMix, the P5_PR1 primer, and an array-specific NGS_R1 primer, run for 10 cycles (98°C/65°C/72°C). Products were cleaned with 1.5× SPRIselect beads and eluted in Buffer EB. Eleven microliters of each cleaned product were then used in a second, nested PCR for another 10 cycles, followed by a second 1.5× SPRIselect cleanup. A final indexing PCR added a unique sample index from the Chromium Dual Index kit (9 cycles), and the indexed products were pooled by accounting for cell number, expected reads, and index sequence, then purified with two rounds of 0.8× SPRIselect cleanup. Final libraries were sequenced on an Illumina NextSeq 2000 using a 500-cycle paired-end v2 kit (28 cycles read 1, 10 cycles I1, 10 cycles I2 index, 350 cycles read 2).

### RNAscope staining

RNAscope in situ hybridization was performed as previously described^32^, using the Multiplex Fluorescent Reagent Kit v2 (Advanced Cell Diagnostics, 323270) according to the manufacturer’s protocol. Tissue sections were first fixed in 4% paraformaldehyde and then progressively dehydrated through a graded ethanol series. Sections were subsequently treated with hydrogen peroxide for 10 min and protease IV for 15 min to permeabilize the tissue and quench endogenous peroxidase activity. Target hybridization was carried out using gene-specific Z probes for 2 h at 40°C in a HybEZ II Hybridization System. Probe signals were then amplified and visualized with TSA Vivid Fluorophores. Nuclei were counterstained with DAPI, and slides were mounted with ProLong Gold Antifade Mountant (Invitrogen, P36930) prior to confocal imaging on a Leica TCS SP8 microscope. The probe used in this study was Mm-Lrrn4-C1 (Cat. No. 443951).

### Lrrn4-CreER mouse generation

An sgRNA (5′-GGTTTGTTTAATCTGGTTGCTGG-3′) was designed to insert a viral P2A self-cleaving peptide sequence and a Cre/ERT2 fusion gene (encoding Cre recombinase fused to the ligand-binding domain of the human estrogen receptor) at the 3′ end of the last exon of the leucine-rich repeat neuronal 4 (Lrrn4) gene. The sgRNA, Cas9 mRNA, and a donor plasmid were microinjected into the cytoplasm of C57BL/6N-derived fertilized eggs with clearly recognizable pronuclei. Correctly targeted embryos were then transferred into pseudopregnant females. Lrrn4-CreER founder mice were identified by PCR and confirmed by Sanger sequencing. F1 male and female mice were subsequently bred to generate homozygous offspring.

### IF staining

Immunofluorescence staining was carried out as described previously^14^. Specifically, hearts were harvested and fixed overnight in 4% paraformaldehyde prior to OCT embedding, after which 10-μm sections were cut and rinsed briefly in PBS to remove residual OCT. Tissue sections were blocked for 1 h at room temperature in a blocking buffer containing 10% goat serum, 1% BSA, and 0.1% Tween 20, then incubated overnight at 4°C with primary antibodies diluted in 1% BSA/PBST. Sections were subsequently washed and probed with the appropriate fluorophore-conjugated secondary antibodies, diluted in blocking buffer, for 1 h at room temperature. After DAPI counterstaining, slides were mounted in Fluoromount-G and imaged by confocal microscopy. Primary antibodies included anti-ESR (Abcam, AB16660), anti-CD31 (BD Biosciences, 550274), anti-Wt1 (Abcam, AB89901), anti-Pdgfra (R&D Systems, AF1062-SP), anti-cTNT (Invitrogen, MA5-12960), and anti-pHH3-488 (Abcam, AB197502). Secondary antibodies were goat anti-rabbit IgG-488 (Thermo Fisher Scientific, A-11008), goat anti-rabbit IgG-647 (Thermo Fisher Scientific, A21244), Goat anti-rat 488 (Thermo Fisher Scientific, A11006), Donkey anti-goat 647 (Thermo Fisher Scientific, A21447), Goat anti-mouse 594 (Thermo Fisher Scientific, A11032).

### Data analysis

#### DARLIN barcode calling

Paired-end sequencing reads were processed with the snakemake_DARLIN workflow (https://github.com/ShouWenWang-Lab/snakemake_DARLIN)^19,20^. Reads were grouped by cell barcode and UMI, and insertion/deletion alleles were called independently at each of the three DARLIN arrays (CA, TA and RA).

#### Clone calling and lineage assignment

Allele tables were converted into clonal identities using MosaicLineage (github.com/ShouWenWang-Lab/MosaicLineage). To suppress homoplasy, each allele was scored against a reference allele bank, and only rare alleles were retained as lineage-informative. A joint clone was defined for each cell by combining its trusted edits across all three arrays (CA, TA, and RA). Array records supporting a joint clone with at least two arrays (joint_allele_num ≥ 2) and matching a cell in the transcriptome dataset were carried forward for clonal analyses (E14.5: 7,506 cells; E18.5: 10,454 cells).

#### Single-cell RNA-seq preprocessing

Transcriptomes captured alongside the barcodes were analyzed in Scanpy^33^ with helper functions from CoSpar (v0.3.3; Python 3.9)^34^. Each timepoint was processed independently from Cell Ranger result. Cells were further filtered to ≥500 detected genes and ≥1,000 total UMIs, and genes to detection in ≥2 cells. Cells were also required to have <10% mitochondrial counts. Highly variable genes were selected on library-size-normalized counts (10,000 counts/cell; min_counts=3, min_cells=3, v-score percentile 85), giving around 3,150 genes. Genes correlated Pearson |r| > 0.25 with a curated cell-cycle panel (Ube2c, Hmgb2, Hmgn2, Tuba1b, Ccnb1, Tubb5, Top2a, Tubb4b, Cdca8, Ccnb2, Mki67, Cdk1) were then removed, leaving E14.5 3,034 and E18 3,036 variable genes.

PCA was computed on the variable genes (40 components, n_neighbors=20, min_dist=0.3). Doublets were scored with Scrublet^35^ and cells with a doublet score >0.3 were discarded. Leiden clustering was used for annotation^36^. Clusters that did not confidently match a fate were left unannotated and excluded from the clonal analyses (E14.5: 18,662/22,372; E18.5:13,046/18,858).

#### Integration of clones with the transcriptome

Joint clone tables were matched to transcriptome barcodes by cell identifier, retaining only lineage-informative cells (joint_allele_num ≥ 2) present in the annotated AnnData object. A clone-by-cell membership matrix (X_clone) was constructed with CoSpar (cs.pp.get_X_clone) and stored on the object (E14.5: 4,048 cells × 2,787 clones; E18.5: 5,042 cells × 4,187 clones).

#### Phylogeny tree diagram

A character-based phylogeny was reconstructed from the DARLIN edits with Cassiopeia^37^. Each joint clone was treated as a leaf and encoded as a binary character vector: every unique locus-tagged edit observed across the CA, TA and RA arrays became one character. Leaves were restricted to informative clones with select_clones: at least 3 cells and at least 2 distinct fates. For E14.5 this yielded 80 clones spanning 607 unique edits, of which 508 were private to a single clone and 99 shared across ≥2 clones; the same criteria were applied to E18.5. Trees were solved with Cassiopeia’s VanillaGreedySolver on the prior-weighted character matrix, and mutationless (zero-edit) edges were collapsed so that internal nodes correspond to genuine branch points. The resulting cladogram was drawn with leaves colored by dominant fate.

#### scMultiome and scATAC-seq data analysis

The embryonic heart single-cell Multiome ATAC + Gene Expression dataset at E14.5 was downloaded from GEO (accession number: GSE272154)^23^ and analyzed using the Seurat V5 pipeline^38^. An independent scATAC-seq dataset^39^ (GEO accession: GSE153479) was also downloaded and processed using the same pipeline.

#### Statistical analysis

Statistical significance between two conditions was assessed using an unpaired, two-tailed Student’s t-test assuming equal variances. Data are presented as mean ± SD. A p-value < 0.05 was considered statistically significant, denoted as *p < 0.05, **p < 0.01, ***p < 0.001, ****p < 0.0001; ns, not significant.

## Acknowledgments

We are grateful to former members of the Li lab, Yuanhang He and Junqi Hu, for their contributions to several experiments, including IF and RNAscope staining analyses. We also thank the Center for Biologic Imaging at the University of Pittsburgh for their support with sample imaging. Computational resources were provided in part by the University of Pittsburgh Center for Research Computing (RRID:SCR_022735), including access to the HTC cluster, which is supported by NIH award S10OD028483. This work was funded by the NIH (R00HL133472 and DP2HL163745 to G.L.) and by an SVRF grant from Additional Ventures awarded to G.L.

## Author contributions

Conceptualization: G.L.

Methodology: P.G., Z.G., H.H., J.X.

Investigation: P.G., Z.G., H.H.

Funding acquisition: G.L.

Project administration: G.L.

Supervision: G.L.

Writing – original draft: G.L.

Writing – review & editing: P.G., Z.G., H.H., J.X., G.L.

## Supplemental Figures

Fig. S1. Lineage tracing with DARLIN system. (A) Diagram of the genetic elements in DARLIN system. (B) The allele frequencies on each targeting arrays including CA, TA, and RA at E14.5 hearts.

Fig. S2. Lineage analysis of mouse embryonic heart development using DARLIN at E18.5. (A) Diagram of the lineage tracing experiment with Dox administration at E10.5 and heart collection at E18.5. (B) Heatmap of the lineage genes expression across the cell types. (C) UMAP plot of the scRNA-seq data labeled with cell lineage information. (D) The number of cells recovered at E18.5 scRNA-seq dataset. (E) The UMI factions on each target sites. (F) Heatmap showing the cellular compositions within each clone in the E18.5 sample. (G) Histogram showing the number of clones at each clone size. (H) Phylogeny of the clones at E18.5. (I) UMAP plots of one representative clone.

Fig. S3. Chromatin accessibility at the Lrrn4 genomic locus. (A) scATAC-seq at P4 showing epicardial-specific chromatin accessibility.

Fig. S4. Heatmap showing the single cell expression pattern of Wt1, Tbx18, Krt19, and Lrrn4 in epicardial cells at three stages.

Fig. S5. Simultaneous RNA staining of Lrrn4 with Tbx18 and Wt1. (A, B) Co-staining at E14.5. (C) Co-staining at P3. Scale bar = 500 μm.

Fig. S6. cTNT staining in ex vivo cultured cardiac cells. eGFP and cTNT staining in cultured cells with or without FACS enrichment. Scale bar = 100 μm.

Fig. S7. Lineage tracing analysis of Lrrn4-CreER–labeled cells. (A) Tamoxifen was given to the Lrrn4-CreER; Rosa26-mTmG pregnant mice at E10.5 and E11.5. (B) Lineage genes for different cell types were stained and imaged together with eGFP. Scale bar = 100 μm.

