## Supplementary figures and images for "Unbiased and Epicardial-Specific Lineage Tracing Reveal Epicardial Contribution to Vascular Endothelial Cells in Heart Development"

### Supplemental figure 1

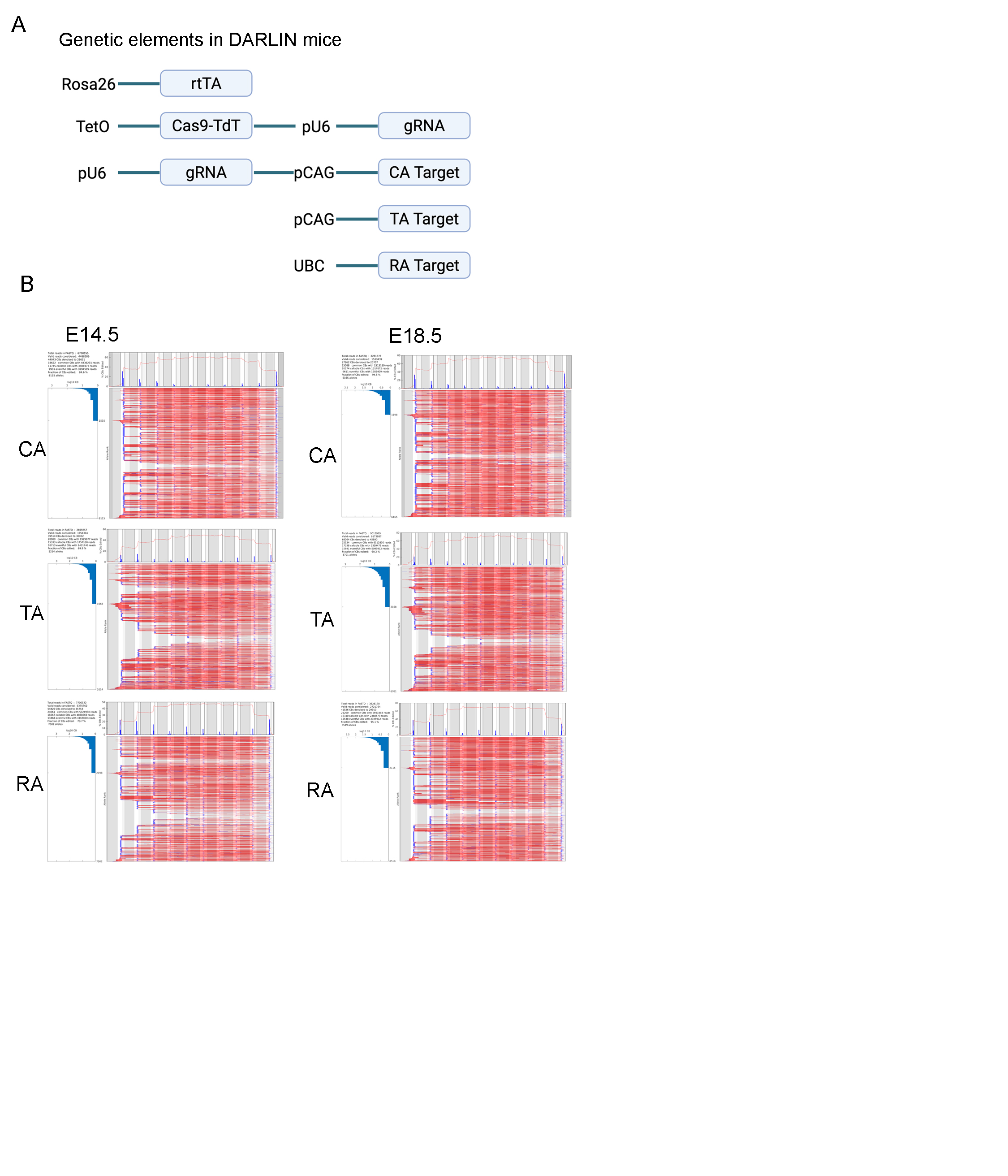

### Supplemental figure 2

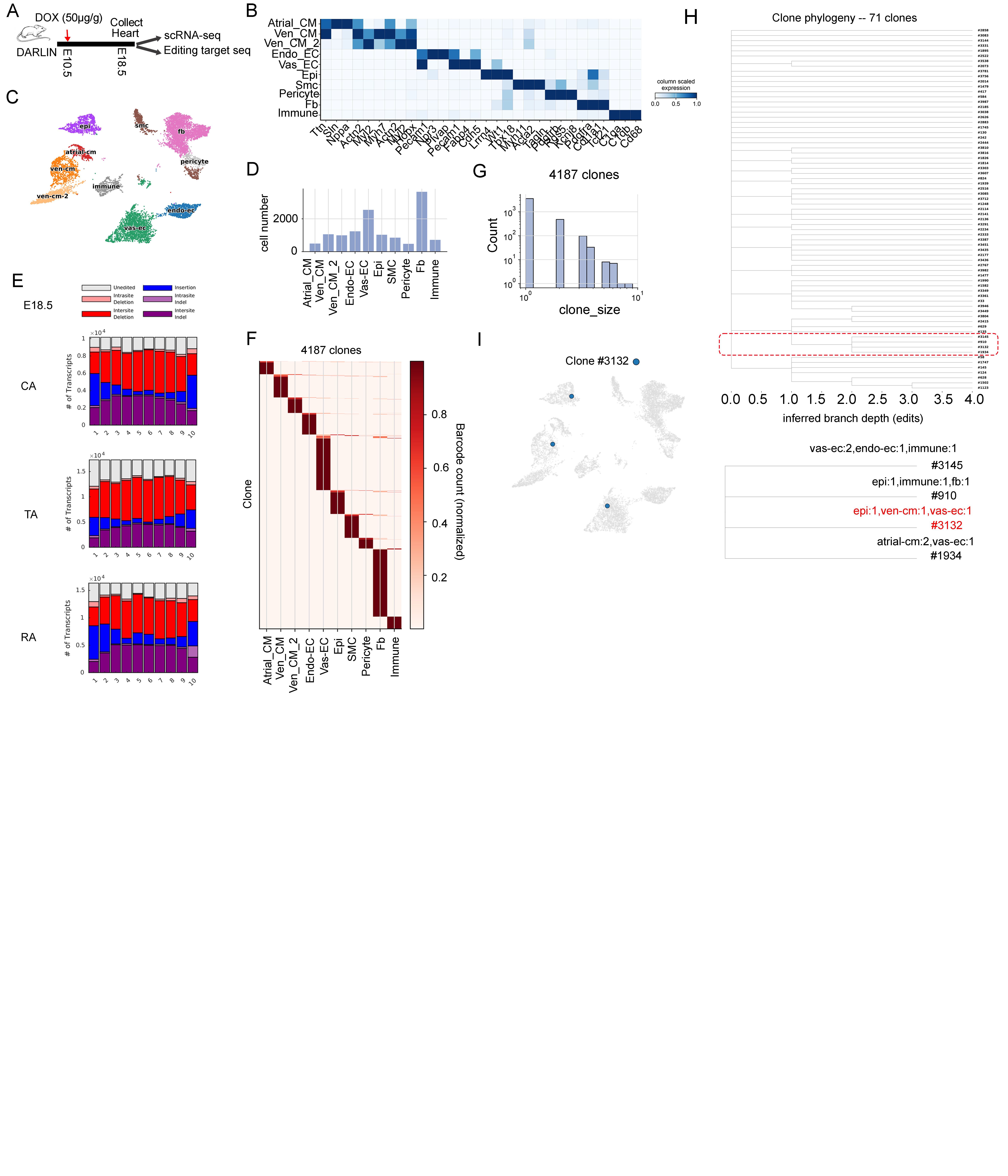

### Supplemental figure 3

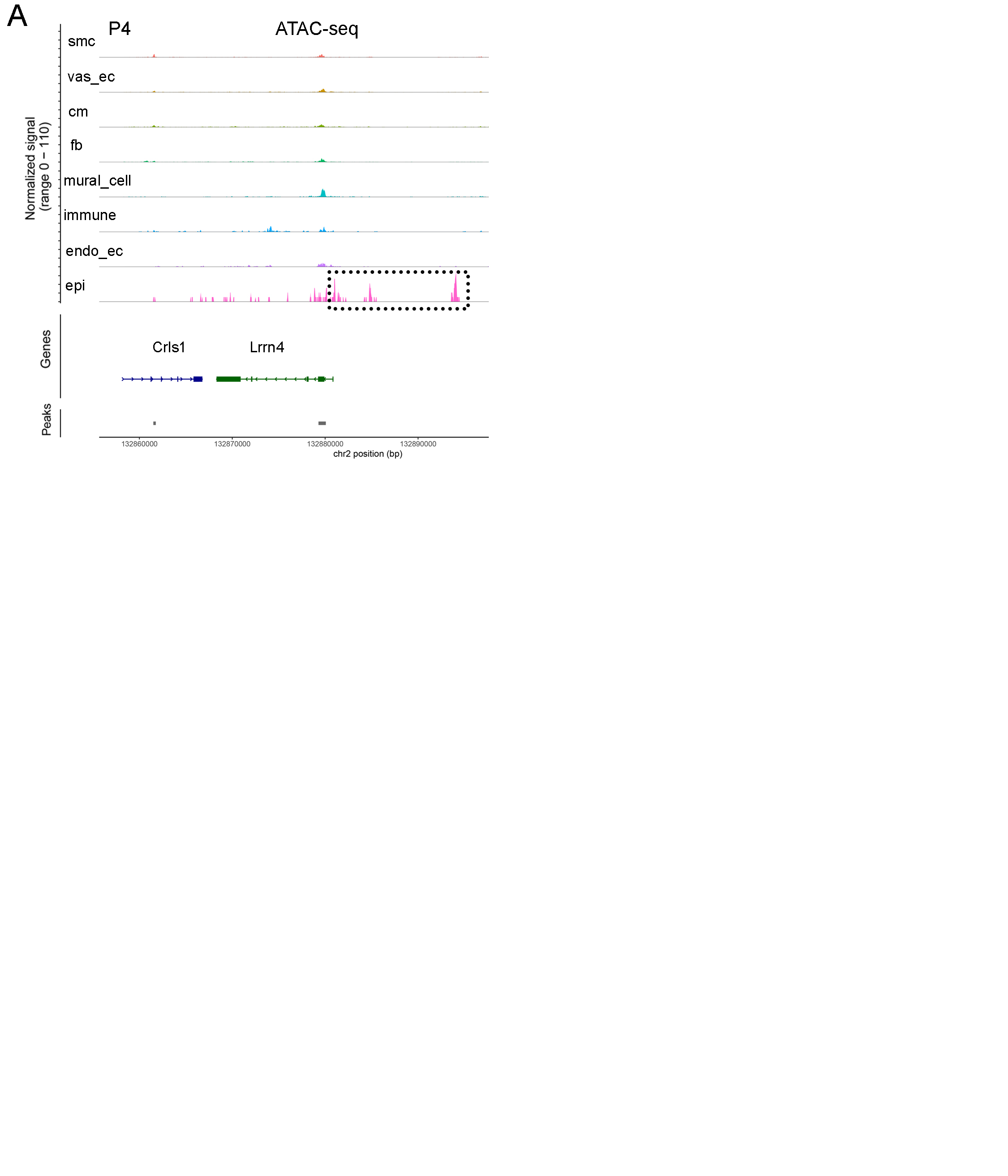

### Supplemental figure 4

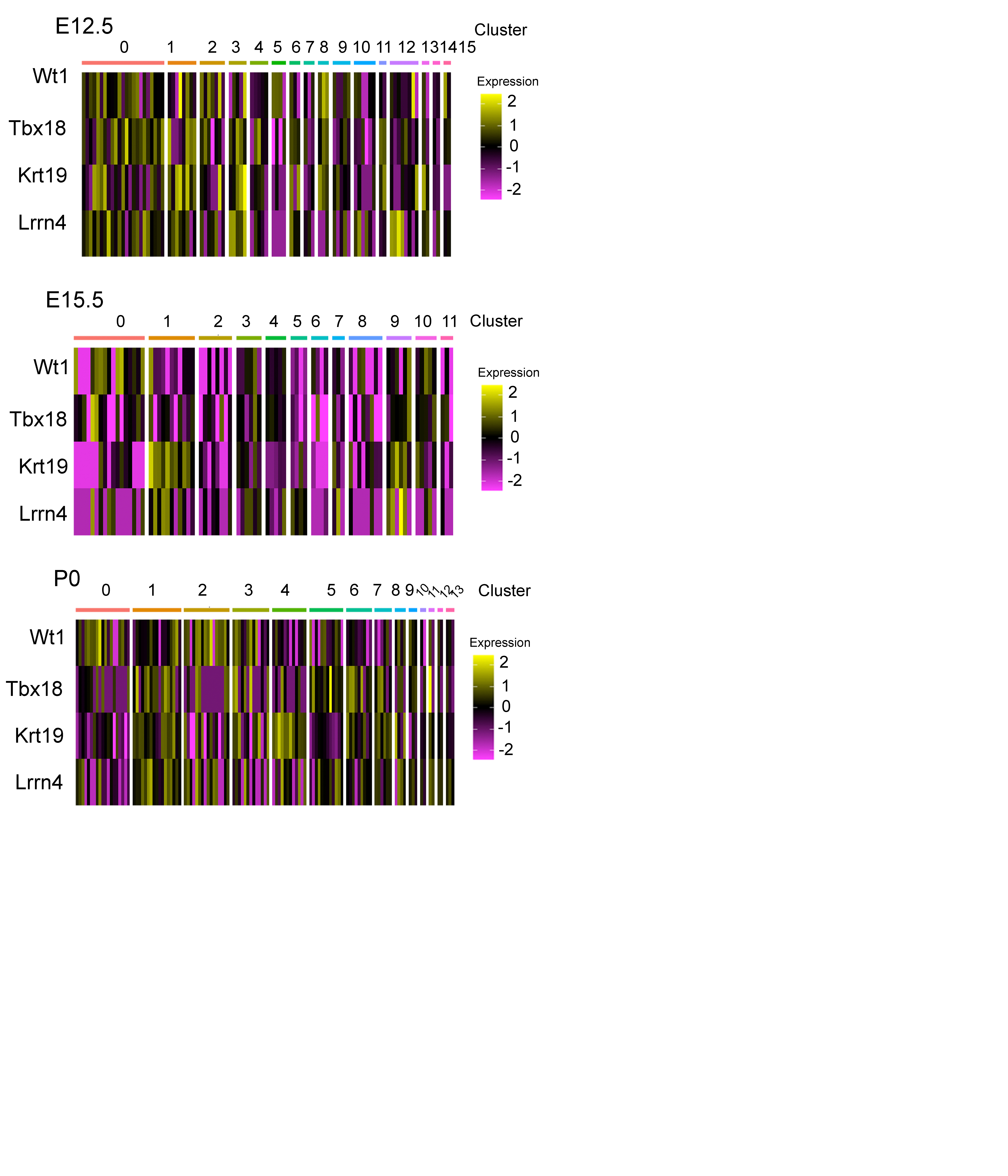

### Supplemental figure 5

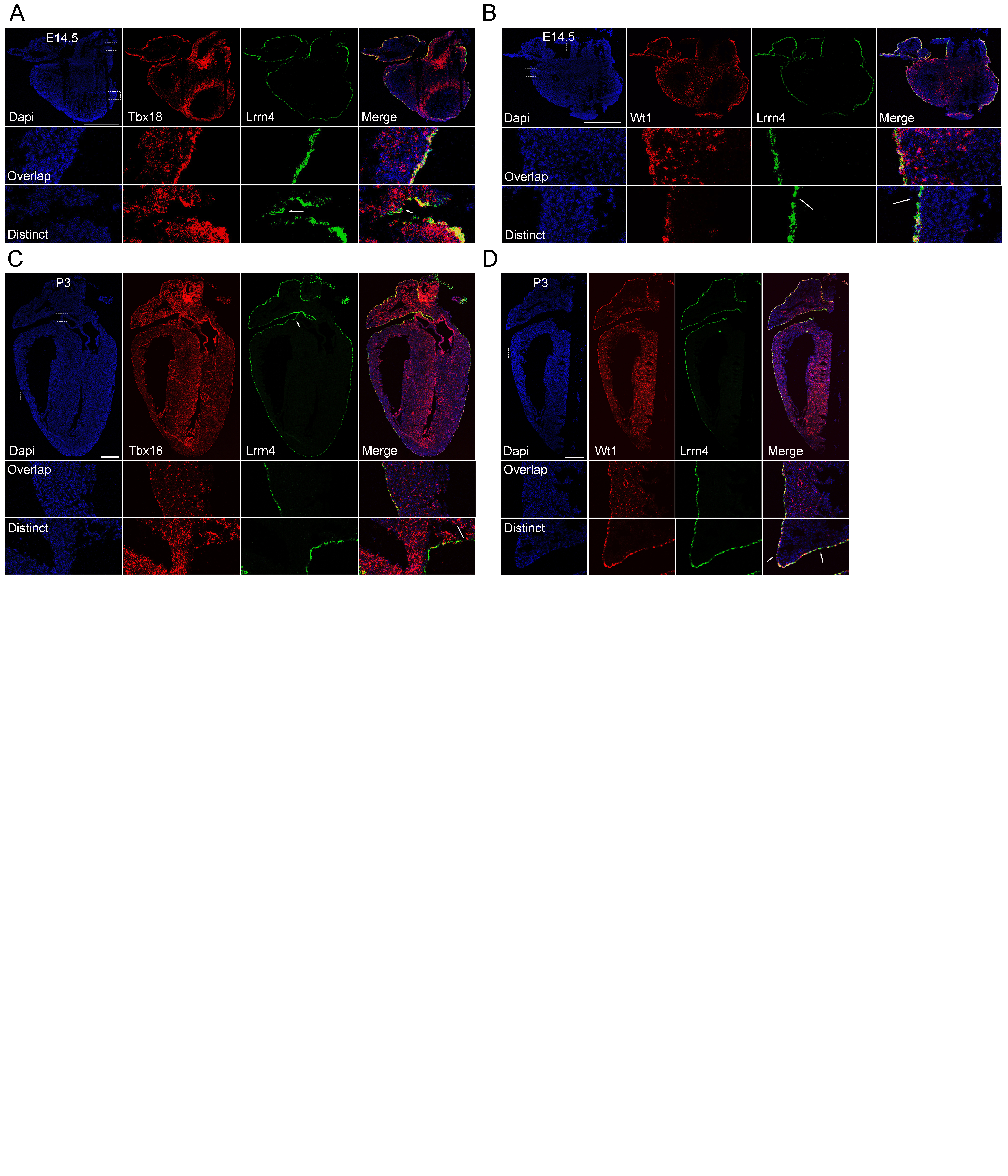

### Supplemental figure 6

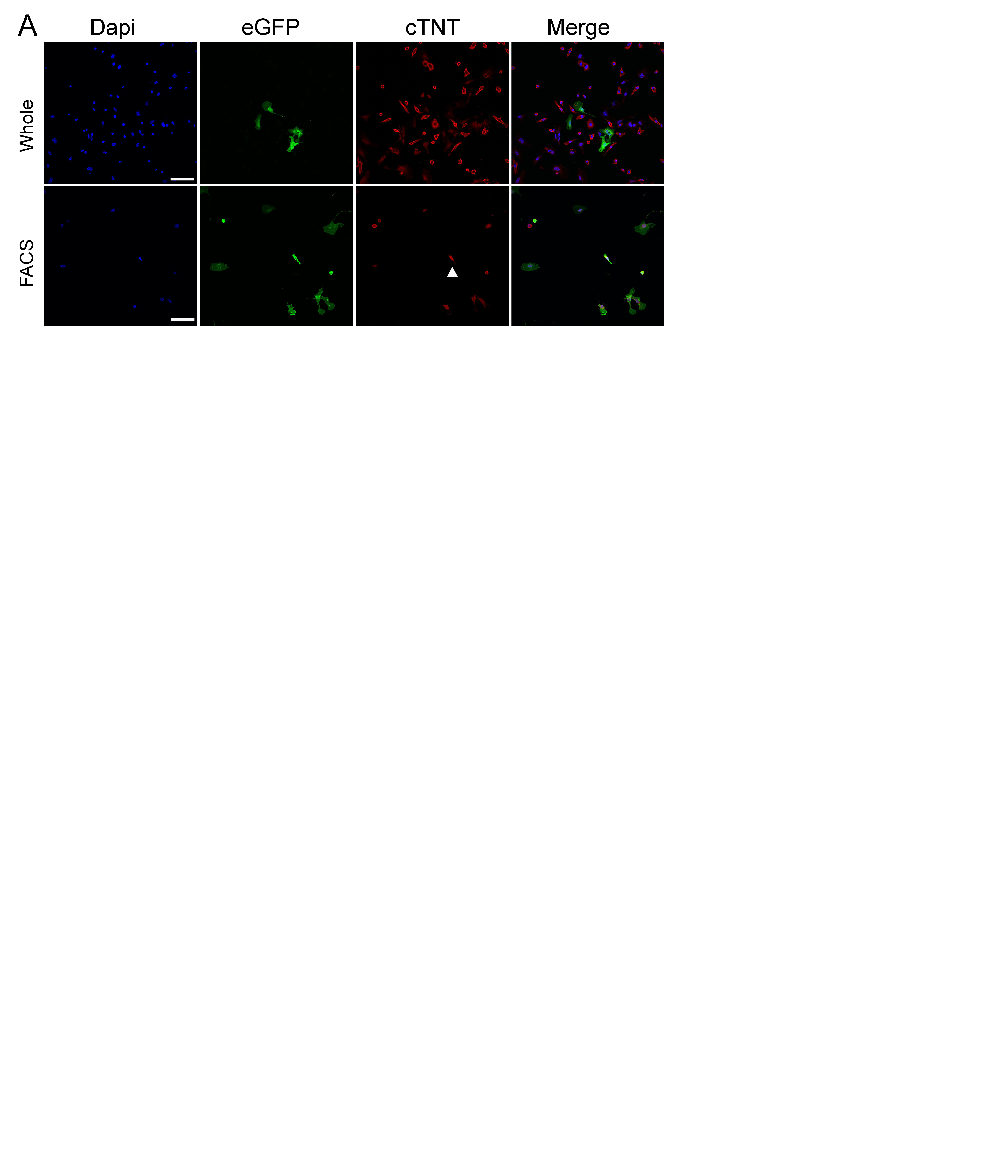

### Supplemental figure 7

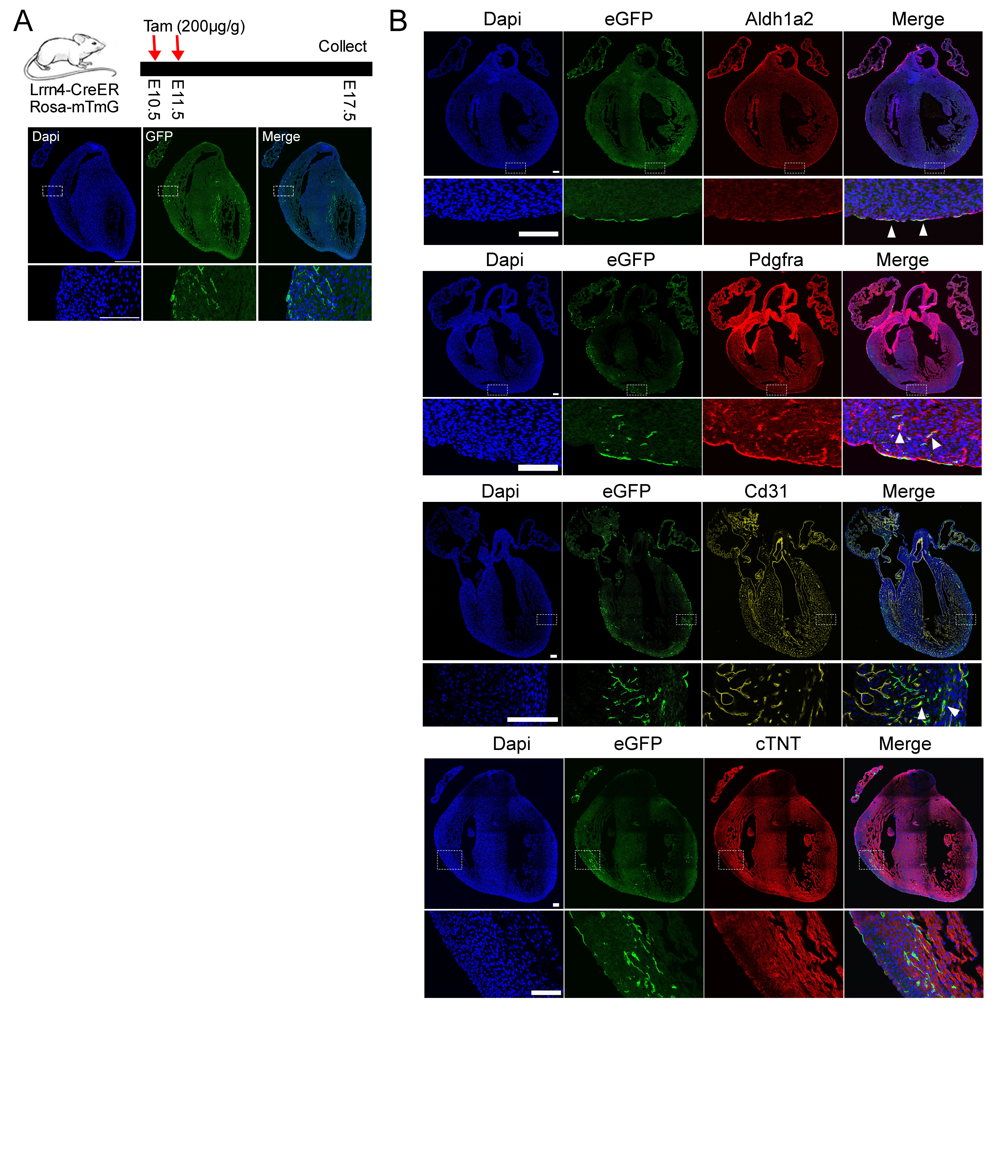
